# GABA_B_ receptors mediate intracellular calcium release in astrocytes of the prefrontal cortex

**DOI:** 10.1101/2024.06.12.598719

**Authors:** Jennifer Bostel, Alina J. Kürten, Antonia Beiersdorfer

## Abstract

The prefrontal cortex (PFC) is a cortical brain region whose multifaceted functions are based on a complex interplay between excitatory pyramidal neurons, inhibitory GABAergic interneurons and astrocytes maintaining a fine-tuned excitation/inhibition balance (E/I balance). The regulation of the E/I balance in cortical network is crucial as the disruption leads to impairments in PFC-associated behavior and pathologies. Astrocytes express specific GABA receptors that mediate intracellular Ca^2+^ signaling upon stimulation by γ-aminobutyric acid (GABA), resulting in the release of gliotransmitters directly impacting information processing. However, the signaling pathway leading to GABA-induced Ca^2+^ signaling in astrocytes of the PFC is not well understood. Here we took advantage of GLAST-promoter driven GCaMP6s expression in astrocytes to study GABAergic Ca^2+^ signaling in PFC astrocytes by confocal microscopy. The results show that GABA induces Ca^2+^ signaling via the stimulation of the metabotropic GABA_B_ receptor in astrocytes. GABA_B_ receptor-mediated Ca^2+^ signals greatly depend on intracellular Ca^2+^ stores rather than on extracellular Ca^2+^. Additionally, antagonists of the PLC/IP_3_-signaling cascade significantly reduced GABA_B_ receptor-mediated Ca^2+^ signaling in astrocytes, suggesting that astrocytic GABA_B_ receptors in the PFC are coupled to the G_q_-GPCR signaling pathway.

## Introduction

Gamma-aminobutyric acid (GABA) serves as the major inhibitory neurotransmitter in the central nervous system, whereas glutamate is considered to be the main excitatory neurotransmitter in vertebrates (Yoon et al., 2012). In general, there are two major classes of neurons in the prefrontal cortex (PFC), approximately 80-90% excitatory glutamatergic pyramidal neurons and 10-20% GABA-releasing interneurons establishing a well-defined neuronal circuit maintaining a fine-tuned balance between excitation and inhibition (E/I balance) (Ferguson & Gao, 2018; Isaacson & Scanziani, 2011; Xu et al., 2019). The E/I balance in neuronal networks is crucial for information processing as the disruption of the E/I balance induces impairments of PFC-associated behavior, such as working memory, social interaction and emotional regulation (Ferguson & Gao, 2018). Additionally, an imbalanced E/I ratio is associated with pathologies of the PFC, such as schizophrenia and autism spectrum disorder (Ko et al., 2015; Pizzarelli & Cherubini, 2011; Sprekeler, 2017). However, not only neurons are involved in the establishment and maintenance of cellular information processing in the CNS, also astrocytes fulfill diverse functions contributing to healthy brain physiology. To do so astrocytes contribute to and regulate the blood brain barrier, supply neurons with metabolites and modulate synaptic transmission in the neuronal network (Allen & Barres, 2009; Volterra & Meldolesi, 2005). In fact, astrocytes have been shown to sense as well as release GABA, thereby modulating synaptic transmission, making astrocytes an ideal cellular partner to contribute to the maintenance and tuning of the E/I balance in cortical networks (Kozlov et al., 2006; Lee et al., 2010; Liu et al., 2000; Mederos et al., 2021; Perea et al., 2016; Yoon et al., 2012). To sense GABA release by neurons, astrocytes express both types of GABA receptors, i.e. the ionotropic GABA_A_ receptor as well as the metabotropic GABA_B_ receptor (Kettenmann et al., 1987; Mederos et al., 2021; Meier et al., 2008). Additionally, GABA transporters, such as GAT1 and GAT3 are present in astrocytes of different brain regions (Boddum et al., 2016; De Biasi et al., 1998; Doengi et al., 2009; Shigetomi et al., 2011). GABA_A_ receptors are chloride channels mediating a chloride influx upon GABA stimulation initiating a hyperpolarization of the cell. GABA_B_ receptors on the other hand are G_i/o_-coupled metabotropic receptors leading to an inhibition of the adenylyl cyclase and the reduction of intracellular cAMP levels. GABA-mediated Ca^2+^ signaling in astrocytes has also been reported in multiple studies (Doengi et al., 2009; Mariotti et al., 2016; Meier et al., 2008), however the exact signaling pathways remain barely understood (Ishibashi et al., 2019). In this study we aimed to investigate the signaling pathway that induces GABAergic Ca^2+^ signaling in astrocytes of the PFC by confocal Ca^2+^ imaging. The results show that GABA evoked GABA_B_ receptor-mediated Ca^2+^ signals in PFC astrocytes that depend on intracellular Ca^2+^ stores and the PLC/IP_3_-signaling cascade, indicating that astrocytic GABA_B_ receptors in the PFC are coupled to the G_q_-GPCR signaling pathway.

## 2 Material and methods

### 2.1 Animals and preparation of prefrontal cortex slices

Mice of the GLAST-Cre^ERT2^ x GCaMP6s^fl/fl^ (age: p28-p60) strain (Mori et al., 2006; Madisen et al., 2015) were kept at the institutional animal facility of the University of Hamburg. Animal rearing and all experimental procedures were performed according to the European Union’s and local animal welfare guidelines (GZ G21305/591-00.33; Behörde für Gesundheit und Verbraucherschutz, Hamburg, Germany). For induction of GCaMP6s expression controlled by the GLAST promoter, tamoxifen (Carbolution Chemicals GmbH, St.Ingbert, Germany) was dissolved in ethanol and Mygliol®812 (Sigma Aldrich) and injected intraperitoneally for three consecutive days (starting p21; 100 mg/kg body weight). Animals were analyzed 14-28 days after the first injection. The prefrontal cortex was removed from the opened head in cooled preparation solution (molarities in mM: 83 NaCl, 1 NaH_2_PO_4_x2H_2_O, 26.2 NaHCO_3_, 2.5 KCl, 70 sucrose, 20 D-(+)-glucose, 2.5 MgSO_4_ x7 H_2_O). Standard artificial cerebrospinal fluid (ACSF) for experiments and storage of the preparations consisted of (molarities in mM): 120 NaCl, 2.5 KCl, 1 NaH_2_PO_4_x2H_2_O, 26 NaHCO_3_, 2.8 D-(+)-glucose, 1 MgCl, 2 CaCl_2_. The modified 0 Ca^2+^-ACSF consisted of (molarities in mM): 120 NaCl, 2.5 KCl, 1 NaH_2_PO_4_x2H_2_O, 26 NaHCO_3_, 2.8 D-(+)-glucose, 3 MgCl, 0.5 EGTA. Preparation solution and ACSF were continuously perfused with carbogen (95% O_2_, 5% CO_2_) to maintain the pH of 7.4 and to supply oxygen.

### 2.2 Reagents

The reagents D-2-amino-5-phosphonovaleric acid (D-APV; antagonist of NMDA receptors; #D-145), 2,3-dioxo-6-nitro-1,2,3,4-tetrahydrobenzo[f]quinoxaline-7-sulfonamide (NBQX; antagonist of AMPA/kainate receptors; #N-186), tetrodotoxin (TTX; inhibiting voltage-gated sodium channels; #T-550) and gabazine (Antagonist of GABA_A_ and Glycine receptors, #G-215) were purchased from Alomone labs (Jerusalem, Israel). The compounds (*R,S*)-4-Amino-3-(4-chlorophenyl)butanoic acid (R,S)-Baclofen; GABAB receptor agonist, #ab120149) and cyclopiazonic acid (CPA; Ca^2+^-ATPase inhibitor; #ab120300) were obtained from Abcam (Cambridge, UK). 2-Aminoethoxydiphenylborane (2-APB; IP3 receptor antagonist; #1224), (2*S*)-3-[[(1*S*)-1-(3,4-Dichlorophenyl)ethyl]amino-2-hydroxypropyl](phenylmethyl)phosphinic acid hydrochloride (CGP 55845 hydrochloride; GABAB receptor antagonist; #1248), gamma-Aminobutyric acid (GABA, endogenous agonist of GABA receptors; #0344), and 1-[6-[[(17β)-3-Methoxyestra-1,3,5(10)-trien-17-yl]amino]hexyl]-1*H*-pyrrole-2,5-dione (U 73122; Phospholipase C inhibitor; #1268) were purchased from BioTechne (Wiesbaden, Germany). Stock solutions were prepared according to the manufacturer’s instructions and dissolved in ACSF at the final concentrations immediately prior to the experiment. Agonists were applied for 30 s, whereas antagonists were applied for 10-30 min via the perfusion system.

### 2.3 Confocal Ca^2+^ Imaging, Data analysis and statistics

For Ca^2+^ imaging experiments, slices of the prefrontal cortex (220 μm, coronal) of GLAST-Cre^ERT2^ x GCaMP6s^fl/fl^ mice were used. Slices were prepared using a vibratome (Leica VT1200S). Coronal PFC slices were transferred into the recording chamber, fixed with a platinum grid covered with nylon strings and continuously perfused with ACSF via the perfusion system. Changes in cytosolic Ca^2+^ concentration in astrocytes were detected by the fluorescence of GCaMP6s (excitation: 488 nm; emission: 500-530 nm) using a confocal microscope (eC1, Nikon, Düsseldorf, Germany). Images were acquired at a time rate of one frame every 3 s. To analyze changes in cytosolic Ca^2+^ in single cells, regions of interest (ROIs) were defined using Nikon EZ-C1 3.90 software. Astrocytes were identified by GLAST promoter-driven GCaMP6s expression. The changes in Ca^2+^ were recorded throughout the experiments as relative changes in GCaMP6s fluorescence (ΔF) with respect to the baseline fluorescence, which was normalized to 100%. Quantification of the Ca^2+^ transients were calculated by the amplitude of ΔF. All values are stated as mean values ± standard error of the mean. The number of experiments is given as n = x/y, where x is the number of analyzed cells and y is the number of animals. At least 3 animals were analyzed in all experiments. Statistical significance was estimated by comparing three means using Friedmann ANOVA and the Wilcoxon post-hoc test for paired data sets, and the Mann-Whitney-U test for unpaired data sets. The Grubbs test was used to identify outliers throughout all experiments. The error probability p was * p< 0.05; ** p< 0.01; *** p< 0.001.

## 3 Results

### 3.1 GABA induces Ca^2+^ signals in astrocytes of the prefrontal cortex

To study GABAergic Ca^2+^ signaling in astrocytes of the PFC we took advantage of GLAST-Cre^ERT2^ x GCaMP6s^fl/fl^ mice, in which the genetically encoded Ca^2+^ indicator (GECI) GCaMP6s is expressed under control of the astrocyte-specific GLAST promoter (Mori et al., 2006; Madisen et al., 2015). GCaMP6s is a commonly used GECI suitable for Ca^2+^ imaging in glial cells (Lohr et al., 2021). Bath application of GABA induced Ca^2+^ signaling in astrocytes in L1-3 of the PFC in a dose-dependent manner (Fig. 1A-D). To identify all responding astrocytes in the field of view, we bath-applied norepinephrine (NE, 10 μM, 30 s) triggering α1-receptor-mediated Ca^2+^ signaling in astrocytes after the different doses of GABA (Suppl. Fig. 1). The number of astrocytes that responded to the application of NE was defined as 100% responding astrocytes (Fischer et al., 2021). It should be noted that approximately 80% of all astrocytes undergo Cre-dependent DNA recombination in these mice (Jahn et al., 2018). Bath application of GABA (50 μM, 30 s) resulted in Ca^2+^ transients that amounted to 8.4 ± 1.43 % ΔF in 16.13 % of the astrocytes (Fig. 1D-F). GABA (200 μM) induced Ca^2+^ transients that amounted to 53.02 ± 4.32 % ΔF in an increasing number of 36.93 % of the astrocytes. The application of 500 μM GABA induced Ca^2+^ transients with an amplitude of 91.93 ± 6.15 % ΔF in 39.52 % of the astrocytes, whereas 1 mM GABA induced Ca^2+^ transients with an amplitude of 152.01 ± 13.85 % ΔF in 55.81 % of the astrocytes. The results indicate that about half of the astrocyte population in L1-3 of the PFC that express GCaMP6s respond to the application of GABA with Ca^2+^ transients. For half-maximal activation of GABA receptor-mediated Ca^2+^ transients in our experiments a concentration of 500 μM GABA was sufficient to induce robust Ca^2+^ signals in a reasonable number of astrocytes of the PFC. To elucidate any indirect effects on GABA-induced Ca^2+^ transients in astrocytes we performed experiments in the presence of TTX (0.5 μM), suppressing action potential firing by neurons and thereby reducing neuronal transmitter release. GABA-induced Ca^2+^ transients were significantly reduced to 82.02 ± 5.99% of the control (n=87/3; p=3.21x10^−7^) in the presence of TTX, suggesting neuronal impact on GABA-induced Ca^2+^ transients in astrocytes (Fig. 1 G,H). Additional inhibition of glutamatergic AMPA (NBQX, 10 μM) and NMDA receptors (D-APV, 100 μM) further reduced GABA-induced Ca^2+^ transients significantly (Fig. 1 I,J). GABA-induced Ca^2+^ transients in astrocytes in the presence of glutamatergic inhibition amounted to 96.64 ± 7.71 % of the control, showing a significant reduction in amplitude compared to GABA-induced Ca^2+^ transients in the presence of TTX (n=125/3; p=1.12×10^−4^). To avoid indirect effects on GABA-induced Ca^2+^ signaling, the following experiments were performed in a combination of all three antagonists.

**Figure 1:**
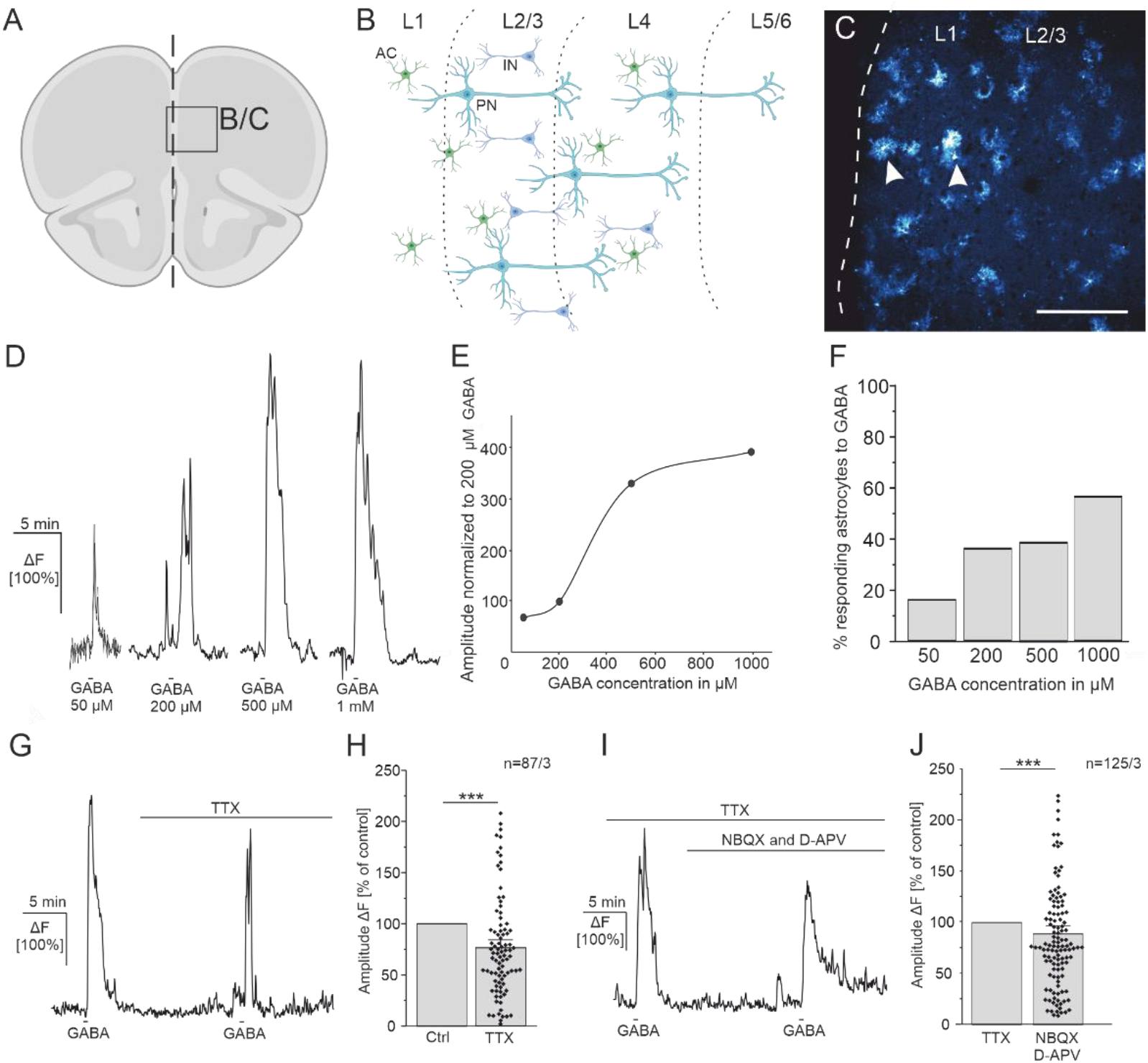
GABA induces Ca^2+^ transients in PFC astrocytes. **(A)** Overview of the prefrontal cortex of mice in coronal orientation. **(B)** Schematic cellular organization of the PFC including pyramidal neurons (PN), interneurons (IN) and astrocytes (AC). Images were created using BioRender. **(C)** GCaMP6s expressing astrocytes in layer 1 and layer 2+3 (L1, L2-L3) of the prefrontal cortex visualized by confocal microscopy. Arrowheads indicate astrocytes that express GCaMP6s controlled by the GLAST promoter. Scale bar: 200 μm. **(D)** Representative traces of Ca^2+^ signaling in astrocytes induced by different doses of GABA. **(E)** Average amplitudes of GABA-induced Ca^2+^ transients normalized to the 200 μM GABA response, showing the dose-response relationship. **(F)** Percentage of astrocytes that respond to different concentrations of GABA with Ca^2+^ transients in comparison to NE application. Note that 100% responding astrocytes were identified by NE-application (Suppl. Fig. 1). **(G)** GABA-induced Ca^2+^ transients are significantly reduced in the presence of TTX (0.5 μM). **(H)** Average amplitudes of GABA-induced Ca^2+^ transients in control conditions compared to TTX. **(I)** GABA-induced Ca^2+^ transients are affected by glutamatergic antagonists (D-APV, 100 μM; NBQX, 10 μM). **(J)** Average amplitudes of GABA-induced Ca^2+^ transients under control conditions compared to glutamatergic inhibitors. *** p<0.001.

### 3.2 GABA_B_ receptors mediate Ca^2+^ transients in astrocytes

Various signaling cascades have been reported leading to GABA-mediated Ca^2+^ signals in astrocytes (Ishibashi et al., 2019). Both, activation of the ionotropic GABA_A_ receptor as well as the metabotropic GABA_B_ receptor lead to Ca^2+^ elevations in culture and in astrocytes of the somatosensory cortex and hippocampus (Mariotti et al., 2016; Meier et al., 2008; Nilsson et al., 1993). Additionally, GABAergic Ca^2+^ transients in developing olfactory bulb astrocytes depend on the reduced Na^+^/Ca^2+^ exchange induced by loading astrocytes with Na^+^ upon GABA uptake (Doengi et al., 2009). To identify the receptors involved in GABA-mediated Ca^2+^ signaling in PFC astrocytes, we tested the effect of the GABA_A_ receptor antagonist, Gabazine (5 μM) and the GABA_B_ receptor antagonist, CGP 55845 (10 μM) on GABA-induced Ca^2+^ responses in astrocytes. In the presence of Gabazine, the amplitude of GABA-induced Ca^2+^ responses did not show any difference and amounted to 106.05 ± 16.22 % of the control (n=23/3; p=0.50) (Fig. 2 A,B). The GABA_B_ receptor antagonist, CGP 55845 on the other hand nearly completely abolished GABA-induced Ca^2+^ transients in astrocytes. The average amplitude of GABA-induced Ca^2+^ transients in the presence of CGP 55845 amounted to 17.90 ± 4.09 % of the control (n=41/3; p=1.11×10^−14^). The results show that GABA-induced Ca^2+^ transients in astrocytes of the PFC are mediated by GABA_B_ receptors (Fig. 2 C,D).

**Figure 2:**
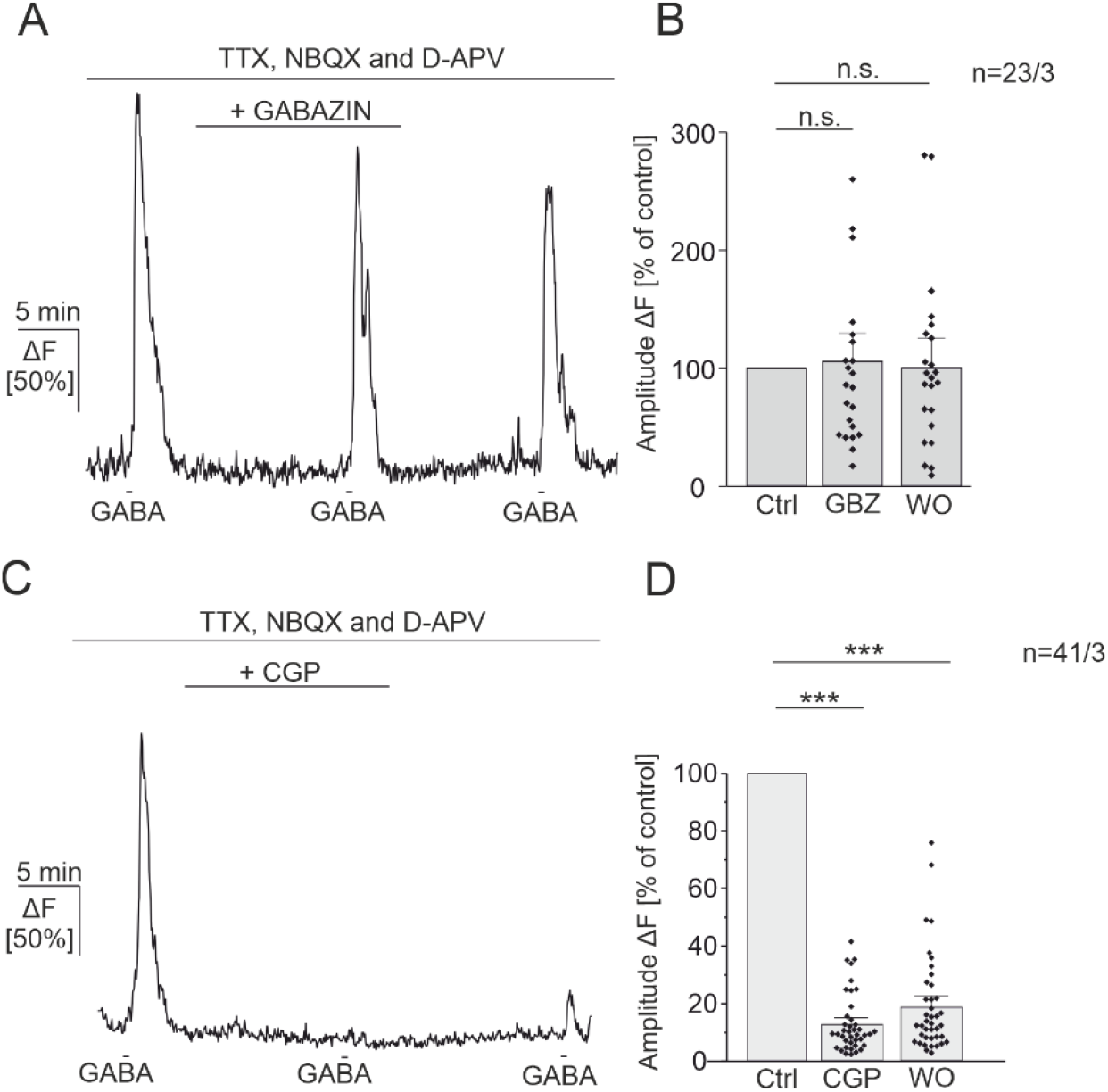
GABA-induced Ca^2+^ transients in astrocytes are mediated by GABA_B_ receptors. **(A)** GABA-induced Ca^2+^ transients are not affected by the GABA_A_ receptor antagonist, Gabazine (5 μM). All experiments were performed in the presence of TTX (0.5 μM), NBQX (10 μM) and D-APV (100 μM) to isolate direct GABAergic Ca^2+^ responses in astrocytes. **(B)** Average amplitudes of GABA-induced Ca^2+^ transients normalized to the control application of GABA, showing no significant difference between control (Ctrl) and gabazin (GBZ, 5 μM). WO: wash out. **(C)** GABA-induced Ca^2+^ transients are strongly inhibited by the GABA_B_ receptor antagonist, CGP 55845 (CGP, 10 μM). **(D)** Average amplitudes of GABA-induced Ca^2+^ transients normalized to the control application of GABA, showing a significant difference in the presence of the GABA_B_-receptor inhibitor. n.s: not significant; *** p<0.001.

### 3.3 GABA_B_-receptor mediated Ca^2+^ transients in astrocytes depend on the PLC/IP_3_-mediated signaling pathway

Usually GABA_B_ receptors are G_i/o_-GPCR-coupled, inducing slow inhibitory signaling pathways in neurons. Presynaptically, GABA_B_ receptor-mediated inhibition of voltage-gated Ca^2+^ channels reduces neurotransmitter release, whereas postsynaptically GABA_B_ receptor-activation recruits inwardly rectifying potassium channels resulting in a slow hyperpolarization of the cell. Contrary, in astrocytes GABA_B_ receptor-activation initiates Ca^2+^ responses, followed by diverse modification mechanisms of synaptic transmission, suggesting the involvement of the G_q_-GPCR-signaling cascade (Kang et al., 1998; Mariotti et al., 2016; Mederos et al., 2021; Perea et al., 2016; Serrano et al., 2006). We aimed to investigate the intracellular signaling pathway that leads to GABA_B_ receptor-mediated Ca^2+^ transients in astrocytes of the PFC. Therefore, we applied the specific GABA_B_ receptor agonist, baclofen to induce robust GABA_B_ receptor-specific Ca^2+^ transients. Bath application of baclofen (200 μM) evoked Ca^2+^ transients in astrocytes with an average amplitude of 220.72 ± 11.77% ΔF. We performed multiple applications of baclofen with a 10-min interval in-between to evaluate a possible decrease in baclofen-induced Ca^2+^ transients by receptor or signaling cascade desensitization (rundown). The second application of baclofen induced Ca^2+^ transients that amounted to 59.70 ± 3.02% of the first baclofen application (control) and showed a significant reduction (n=162/3; p=0.00, Fig. 3 A,B). Therefore, in the following, baclofen-induced Ca^2+^ transients in astrocytes in presence of different antagonists are compared to the corresponding baclofen application in the rundown experiment. To control the agonist specificity of baclofen, we applied baclofen in the presence of the GABA_B_ receptor antagonist, CGP 55848. In the presence of CGP 55848, baclofen-induced Ca^2+^ transients in astrocytes were highly and significantly reduced and amounted to 10.3 ± 1.11% of the control, showing specific activation of GABA_B_ receptors in astrocytes (p=2.41 x 10^−34^, Fig. 3 C,D).

**Figure 3:**
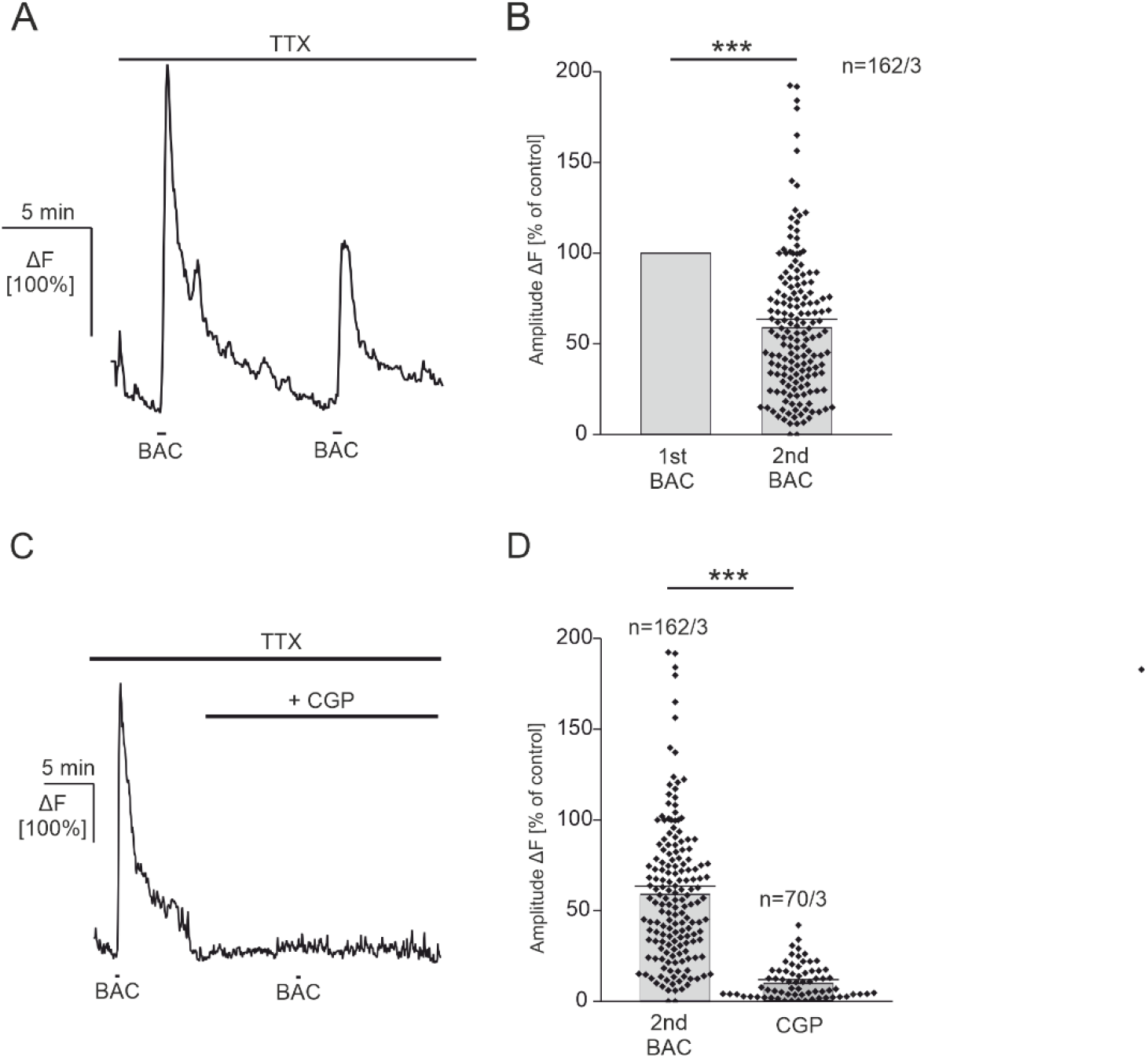
Baclofen-induced Ca^2+^ transients in astrocytes of the PFC. **(A)** Repetitive applications of baclofen (200 μM) result in reduced baclofen-induced Ca^2+^ responses over time (rundown). All experiments were performed in the presence of TTX (0.5 μM). **(B)** Average amplitudes of the 1^st^ and 2^nd^ baclofen-induced Ca^2+^ response (1^st^ and 2^nd^ BAC), showing a significant difference. Therefore, in the following experiments baclofen-induced Ca^2+^ transients under experimental conditions are always compared to the corresponding rundown experiment. **(C)** Baclofen-induced Ca^2+^ transients in astrocytes are strongly reduced in the presence of the GABA_B_-inhibitor, CGP 55845 (CGP, 10 μM). **(D)** Average amplitudes of baclofen-induced Ca^2+^ transients in the presence of CGP 55845 compared to the 2^nd^ baclofen-application in the rundown experiment. *** p<0.001.

To investigate the signaling pathway responsible for GABA_B_ receptor-mediated Ca^2+^ transients in astrocytes, we first removed extracellular Ca^2+^. In Ca^2+^-free ACSF (0 Ca^2+^), baclofen induced Ca^2+^ transients that amounted to 49.90 ± 2.49% of the control and were not significantly different compared to the rundown (p=0.09), suggesting a minor impact of extracellular Ca^2+^ (Fig. 4 A,C). Intracellular Ca^2+^ store depletion by the inhibition of the SERCA pumps by cyclopiazonic acid (CPA, 20 μM), on the other hand, almost completely abolished baclofen-induced Ca^2+^ transients in astrocytes. Baclofen-induced Ca^2+^ transients in the presence of CPA showed reduced amplitudes that amounted to 6.98 ± 0.81% of its control, showing a significant contribution of intracellular Ca^2+^ stores on GABA_B_ receptor-mediated Ca^2+^ signaling (p=1.21 x 10^−32^; Fig. 4 B,C). It should be noted that Ca^2+^ store depletion by SERCA-pump inhibition using CPA itself induces a rise in intracellular Ca^2+^ levels (Fig. 4B, arrow). We further elucidated the involvement of the phospholipase C (PLC) and IP_3_-receptors in the signaling cascade by using specific antagonists. In the presence of the IP_3_-receptor antagonist, 2-APB (100 μM) baclofen-induced Ca^2+^ transients amounted to 43.97 ± 7.15% of the control and were significantly reduced compared to the rundown (p=9.11×10^−5^). In the corresponding rundown experiment, baclofen-induced Ca^2+^ transients amounted to 92.23 ± 7.19% of its control (Fig. 4 D,F). Moreover, the application of the PLC-inhibitor U73122 (50 μM) also reduced baclofen-induced Ca^2+^ transients in astrocytes to 77.64 ± 5.04% of its control (p=0.03) (Fig. 4 E,F). The results show that GABA_B_ receptor-mediated Ca^2+^transients in astrocytes greatly depend on intracellular Ca^2+^ stores triggering the PLC/IP_3_-signaling cascade.

**Figure 4:**
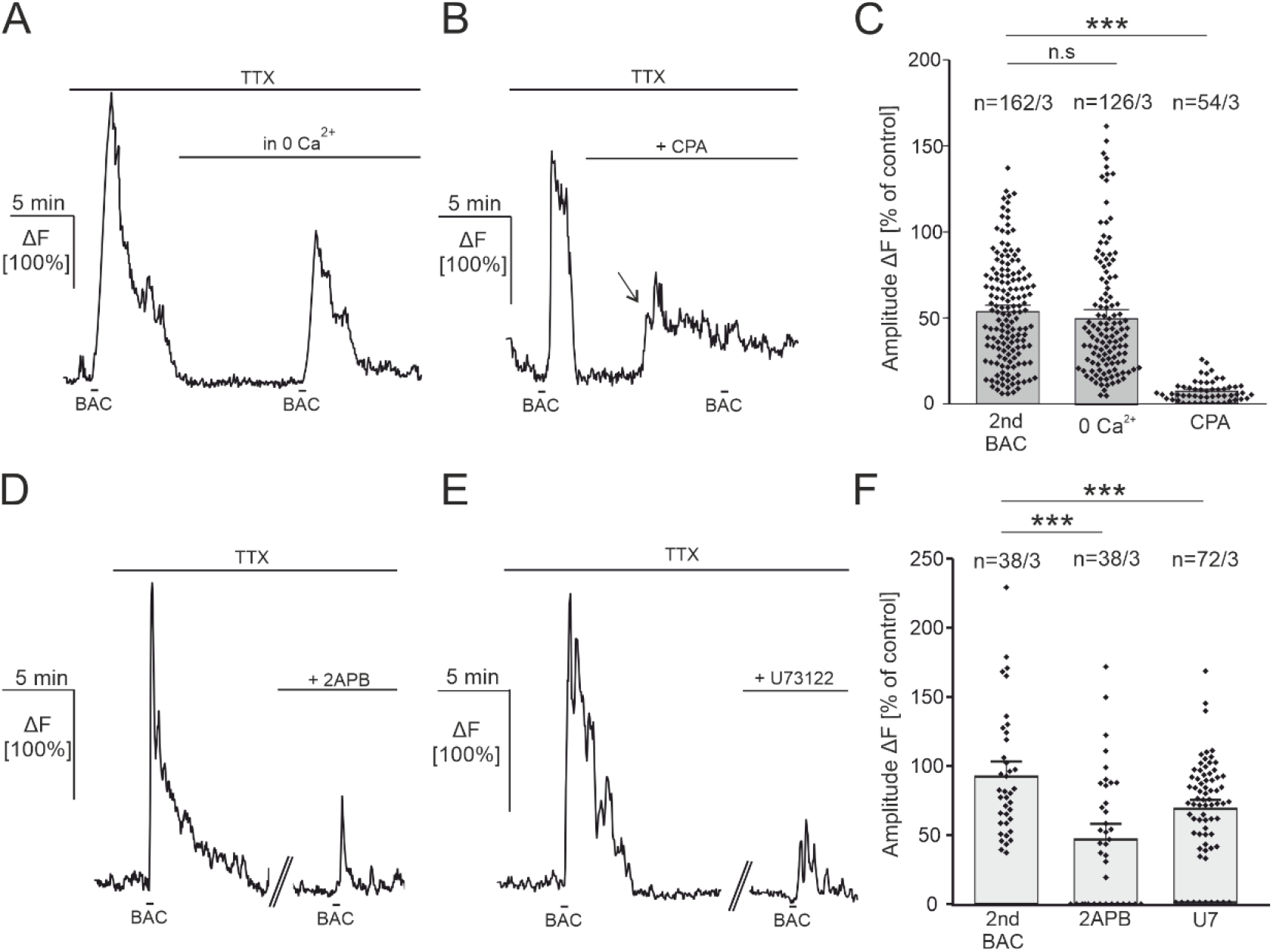
Baclofen-induced Ca^2+^ transients originate from internal Ca^2+^ stores and depend on the PLC/IP_3_ signaling pathway. **(A)** Baclofen-induced Ca^2+^ transients in astrocytes are not affected by the removal of extracellular Ca^2+^ (0 Ca^2+^). **(B)** Baclofen-induced Ca^2+^ transients are abolished in the presence of the SERCA pump inhibitor CPA (20 μM). Application of CPA and the subsequent Ca^2+^ store depletion results in a rise in intracellular Ca^2+^ levels (arrow). **(C)** Average amplitudes of baclofen-induced Ca^2+^ transients in 0 Ca^2+^ and CPA compared to the corresponding rundown experiment. **(D)** The IP_3_-inhibitor, 2-APB (100 μM) reduced baclofen-induced Ca^2+^ transients in astrocytes. **(E)** The PLC-inhibitor, U73122 (U7, 50 μM) reduced baclofen-induced Ca^2+^ transients. **(F)** Average amplitudes of baclofen-induced Ca^2+^ transients in the presence of 2APB and U73122 compared to the corresponding rundown experiment. Note that 2-APB and U73122 were washed in 30 min prior to the 2^nd^ baclofen application, therefore an additional rundown experiment, matching the exact timing of the experiment, was performed for comparison. n.s: not significant, *** p<0.001.

## 4 Discussion

In the present study we investigated GABA-mediated Ca^2+^ signaling in astrocytes of the prefrontal cortex of adult mice. Moreover, we deciphered the intracellular signaling cascade leading to the GABA-induced rise in the intracellular Ca^2+^ concentration in astrocytes.

### 4.1 GABA_B_ receptor-mediated Ca^2+^ signaling in astrocytes depend on the G_q_-protein coupled signaling pathway

In astrocytes the most common Ca^2+^ signaling mechanism is the G_q_-GPCR-coupled pathway, resulting in the IP_3_-dependant Ca^2+^ release from internal stores, just like the endoplasmic reticulum. Classically, neither the ionotropic GABA_A_ receptor, nor the metabotropic GABA_B_ receptor are G_q_-coupled receptors. Whereas GABA_A_ receptors act as chloride channels, usually inducing a hyperpolarization of the cell, the GABA_B_ receptor is G_i/o_-GPCR-coupled reducing the intracellular cAMP concentration by adenylyl cyclase activity reduction. However, astrocytes have been shown to express both GABA-receptor subtypes, both being linked to intracellular Ca^2+^ signaling (Kang et al., 1998; Mederos et al., 2021; Meier et al., 2008; Serrano et al., 2006). In this study we demonstrate that GABA induced Ca^2+^ transients in astrocytes of the PFC, which are not affected by the GABA_A_-receptor antagonist Gabazine but significantly reduced by the GABA_B_-receptor antagonist, CGP 55848. Consequently, GABA_B_ receptors are crucial for GABA-mediated Ca^2+^ signaling in the PFC, as it has been shown for other cortical regions and the developing hippocampus (Mariotti et al., 2016; Meier et al., 2008). However, the molecular pathway leading to functional GABA-mediated Ca^2+^ signaling greatly varies between different brain regions. In contrast to the cortex, in the developing olfactory bulb, astrocytic GABAergic Ca^2+^ signaling mainly relies on GABA-transporter activity (Doengi et al., 2009). Additionally, we show that GABA_B_ receptor-mediated Ca^2+^ transients are diminished after intracellular store depletion by inhibition of the SERCA pumps with CPA, indicating the contribution of intracellular Ca^2+^ stores. Hence, the involvement of the PLC/IP_3_-signaling cascade upstream to the Ca^2+^ release from intracellular Ca^2+^ stores is likely. However, Mariotti et al. (2016) showed that besides the G_q_-GPCR pathway, the inhibition of the G_i/o_-GPCR pathway also greatly reduced GABA_B_-receptor mediated Ca^2+^ oscillations in astrocytes, arguing that the G_q_- and G_i/o_-pathway might be linked for functional GABAergic Ca^2+^ signaling in astrocytes of the cortex. Favoring this hypothesis, it has been shown that chemogenetic G_i/o_-GPCR activation leads to Ca^2+^-signaling in astrocytes of the hippocampus as well (Durkee et al., 2019). However, astrocytic GABA-mediated Ca^2+^ oscillations in the somatosensory cortex are greatly reduced in IP_3_R2 knock-out mice (Mariotti et al., 2016). In line with that, our results show a significant reduction of GABA_B_ receptor-mediated Ca^2+^ signaling after pharmacological inhibition of both the IP_3_-receptor (2APB) and the PLC (U73122), underlining the necessity of the G_q_-GPCR-signaling pathway for GABAergic Ca^2+^ signaling in astrocytes of the PFC.

### 4.2 GABA-mediated Ca^2+^ signaling in astrocytes modulate synaptic transmission in the prefrontal cortex

It is generally accepted that astrocytes sense neuronal activity by a whole repertoire of different neurotransmitter receptors, resulting in functional Ca^2+^ signaling and the release of so-called gliotransmitters, thereby modulating the neuronal network (Allen & Barres, 2009; Araque et al., 1999). As GABA as a neurotransmitter is generally considered to act inhibitory in neurons, we and others have shown that GABA in fact induces “excitatory” Ca^2+^ signals in astrocytes by GABA_B_ receptor activation (Durkee et al., 2019; Mariotti et al., 2016; Meier et al., 2008). As a consequence of astrocytic Ca^2+^ elevations, gliotransmitters are released, acting on neighboring neurons influencing the neuronal network (Araque et al., 1998; Bezzi et al., 1998; Perea & Araque, 2007). Additionally, astrocytic Ca^2+^ signals are crucial to adjust local blood flow to neuronal activity, fueling the cellular network with metabolites (Attwell et al., 2010; Beiersdorfer et al., 2019; Gordon et al., 2007; Mulligan & MacVicar, 2004; Petzold et al., 2008). In the cortex, astrocytes respond to GABA release by different GABAergic interneuron subtypes, such as parvalbumin (PV), somatostatin (SST) and cholecystokinin (CCK) expressing interneurons (Crosby et al., 2018; Deemyad et al., 2018; Mariotti et al., 2018; Mederos & Perea, 2019). Therefore, PV-Interneuron-to-astrocyte communication in the PFC results in GABA_B_ receptor-mediated Ca^2+^ transients in astrocytes and the consequent release of glutamate, thereby modulating the E/I balance of the cortical circuit, directly affecting animal behavior (Mederos et al., 2021; Perea et al., 2016). Additionally, in astrocyte-specific GABA_B_ receptor knock-out mice theta and gamma oscillations in cortical and hippocampal neurons are impaired (Mederos et al., 2021; Perea et al., 2016) Thus, GABA_B_-receptor mediated Ca^2+^ signaling and the involved intracellular signaling cascade is crucial for the modulation of the E/I balance in cortical networks.

## Acknowledgment

We gratefully acknowledge the professional technical assistance of Anne Catrin Rakete, Marion Fink and Steffen Kubitz. We thank Prof. Dr. Frank Kirchhoff and Dr. Anja Scheller for providing transgenic animals.

## Contribution

JB, AJK and AB performed the experiments and analyzed the data. AB designed the experiments and wrote the manuscript. All authors approved the manuscript.

## Data availability statement

All data of this study will be made available by the authors upon request.

