## Supplementary Figure for "GABA_B_ receptors mediate intracellular calcium release in astrocytes of the prefrontal cortex"

### Supplementary material

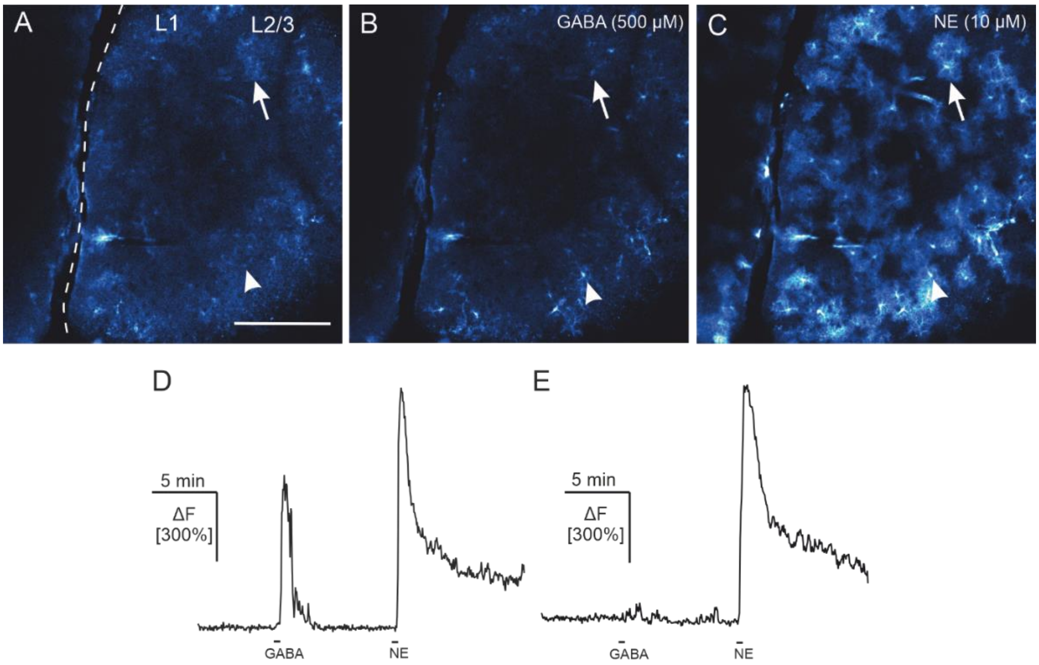


**Suppl. Figure 1: GABA and Norepinephrine (NE) induce $Ca^{2+}$ signaling in PFC astrocytes. (A)** Confocal image of GCaMP6s expression in astrocytes of the PFC. Scale bar: 200 µm. **(B)** Astrocytes respond to GABA application (500 µM) with $Ca^{2+}$ transients. **(C)** Virtually all astrocytes respond to NE application (10 µM) with $Ca^{2+}$ transients. The arrowhead indicates the ROI depicted in D. The arrow indicates the ROI depicted in E. **(D)** An example trace of an astrocyte that responds to GABA and NE. **(E)** An example trace of an astrocyte only responding to NE and not to GABA.
